# Multi-walled carbon nanotubes/nano-Ag-TiO2 membrane DNA electrochemical biosensor

**DOI:** 10.1101/618488

**Authors:** Zhongguo Zheng, Lisa Schultz, John Smith

**Affiliations:** Department of Biochemistry, University of South Carolina, Columbia SC, 29208

## Abstract

A highly sensitive DNA electrochemical biosensor was prepared based on multi-walled carbon nanotube/nano-Ag-TiO2 composite membrane. The Ag-TiO2 composite is mixed with a suitable amount of multi-walled carbon nanotubes (MWNT) dispersed in N,N-dimethylformamide to form a uniform and stable mixed solution, which is applied onto the surface of the bare carbon paste electrode. A MWNT/Ag-TiO2 modified carbon paste electrode was prepared. The large specific surface area and good electron transport properties of carbon nanotubes have a good synergistic effect on the good biocompatibility of Ag-TiO2 nanocomposites and excellent adsorption capacity of DNA, which significantly improves the immobilization and DNA hybridization of DNA probes. Detection sensitivity. The preparation of the sensing membrane and the immobilization and hybridization of DNA were characterized by cyclic voltammetry and electrochemical impedance spectroscopy. The exogenous glufosinate acetyltransferase gene fragment of transgenic plants was detected by electrochemical impedance spectroscopy. The linear range was 1. 0 × 10 - 11 ∼1. 0 × 10 - 6 mol / L. The detection limit was 3. 12 × 10 - 12 mol / L.

## Introduction

Since Iijima discovered carbon nanotubes (CNTs) in 1991, it has been widely used in the sensor field due to its advantages of wide electrochemical window, fast electron transfer rate, good biocompatibility and high mechanical properties [1]. In recent years, CNT-related nanocomposite membranes have attracted much attention because of their synergistic properties [2-5]. Our research team has carried out some research work on the application of CNT-related nanocomposite membranes to prepare DNA electrochemical biosensors [6-8]. Nano-TiO2 is a promising nanomaterial in the field of photochemistry and biochemistry. Its excellent biocompatibility, easy adsorption of biomolecules and good chemical reactivity have been widely used in biosensing [9,10]. Ag nanoparticles are an important part of material synthesis, and the extinction coefficient of biomarker Ag nanoparticles is about 4 times that of gold nanoparticles under the same conditions [11]. The Ag-TiO2 composite combines the advantages of both nano-TiO2 and Ag nanoparticles to attract widespread attention [12,13]. In addition, the electron transfer between Ag and TiO2 also greatly enhances the chemical reactivity of the Ag-TiO2 complex [14,15]. Based on the high sensitivity of multi-walled carbon nanotubes (MWNT) and Ag-TiO2 nanocomposite membranes, a DNA electrochemical biosensor was prepared and the transgenic maize exogenous glufosinate acetyltransferase (PAT) was obtained by electrochemical impedance spectroscopy. Gene fragments are detected. This sensor has good selectivity, stability and regenerability.

## Materials and Method

CHI 660 C Electrochemical Workstation (Shanghai Chenhua Instrument Co., Ltd.), the working electrode is carbon paste electrode or its modified electrode, the counter electrode is platinum wire electrode, the reference electrode is saturated calomel electrode (SCE); pHS-25 pH meter (Shanghai Lei Magnetic Instrument Factory); KQ-50B Ultrasonic Cleaner (Kunshan Ultrasonic Instrument Co., Ltd.); JSM-5900 Scanning Electron Microscope (Japan JEOL Company); Ekpo Ultra Pure Water System (Chongqing Haoyang Enterprise Development Co., Ltd.). Graphite powder (Shanghai colloidal chemical plant, particle size ≤30 μm); High-efficiency slicing paraffin (Shanghai Hualing Rehabilitation Equipment Factory); Ag-TiO2 composite prepared according to the literature [16]; MWNT (Shenzhen Nano Port Co., Ltd., purity > 95) %, diameter < 10 nm, length: 5 to 15 μm); Volume 38, March 2010 Analytical Chemistry (FENXI HUAXUE) Research Report Chinese Journal of Analytical Chemistry No. 3 301 ∼ 306 Detection of exogenous PAT gene fragment material of transgenic maize (SBS Gene Technology Co., Ltd. synthesized, each DNA sequence as described in the literature [17]); DNA fixative with 5. 0 mmol / L pH 7. 0 Tris-HCl buffer solution (containing 50.0 mmol / L NaCl pH 7 0) Prepare and store at 4 °C. The DNA hybridization solution was prepared with a 2 × SSC solution (0.30 mol / L NaCl + 0.03 mol / L sodium citrate). All reagents were of analytical grade and the experimental water was ultrapure water.

The carbon paste electrode and the MWNT/Ag-TiO2 modified carbon paste electrode were prepared by weighing graphite powder 4. 5 g, solid paraffin 1. 5 g, heating and stirring at 80 ° C to make it evenly mixed, and then into a clean glass tube (In the diameter of 4 mm), the copper wire is inserted as a wire, and the carbon paste electrode is prepared by pressing and cooling under a balanced pressure, which is recorded as CPE, and the surface of the electrode is polished to a smoothness on the weighing paper before use. Disperse 1 mg MWNT in 40 mL of 12 mol / L HCl-16 mol / L HNO3 (3:1, V / V) mixed solution, sonicate in water bath for 5 h, filter and wash with ultrapure water until filtration the liquid was made neutral and dried in a vacuum to form a powder for use. 1 mg of the MWNT treated by the above method was taken and dispersed in 1 mL of N,N-dimethylformamide (DMF). 1 mL of MWNT-DMF solution and 1 mg of AgTiO2 complex were added to 50 mL of DMF and sonicated for 5 min to uniformly disperse. A 5 μL drop was applied to the surface of the bare carbon paste electrode and allowed to dry naturally to obtain a MWNT / Ag-TiO2 modified carbon paste electrode. Other modified electrodes are prepared in a similar manner.

Immobilization and hybridization of DNA probe on MWNT/Ag-TiO2/CPE electrode MWNT /Ag-TiO2 /CPE electrode was immersed in 2.0 mL Tris-HCl buffer solution containing 1 μmol / L ssDNA probe and adsorbed at room temperature. After 2 h, it was washed with 0.2% sodium dodecyl sulfate (SDS) solution, and then washed with ultrapure water to remove unfixed ssDNA to obtain ssDNA/MWNT/Ag-TiO2/CPE electrode. The electrode was placed in a hybridization solution containing 1.0 μmol / L of complementary target DNA (cDNA), hybridized at 45 ° C for 60 min, removed, washed with 0.2% SDS solution, and then washed with ultrapure water to remove The hybridized cDNA was obtained as a double-stranded DNA (dsDNA) modified electrode and was designated as dsDNA/MWNT/Ag-TiO2/CPE.

Cyclic voltammetry curves were recorded on CHI 660C in 1.0 mmol/L K4Fe(CN) 6 /K3Fe(CN) 6 (1:1, V /V) 0.1 mol / L KCl solution. The speed is 100 mV/s. The electrochemical impedance spectroscopy curve was also recorded on CHI 660C at room temperature. The detection solution was 1.0 mmol/L K4Fe(CN)6/K3Fe(CN)6 (1:1, V/V). 0. 1 mol / L KCl solution. The applied constant potential is 0. 172 V (v. SCE), and the frequency is 0.1 to 1. 0 × 104 Hz.

## Results and Discussion

### Ag-TiO2 Morphology

Scanning electron microscopy images of Ag-TiO2 nanocomposites (Fig. 1) show that the Ag-TiO2 nanocomposites are composed of many distinct concave spheres with uniform size and smooth surface, with a wall thickness of about 40-80 nm. This structure is beneficial for improving the sensitivity of chemical sensors [18, 19].

**Figure 1.**
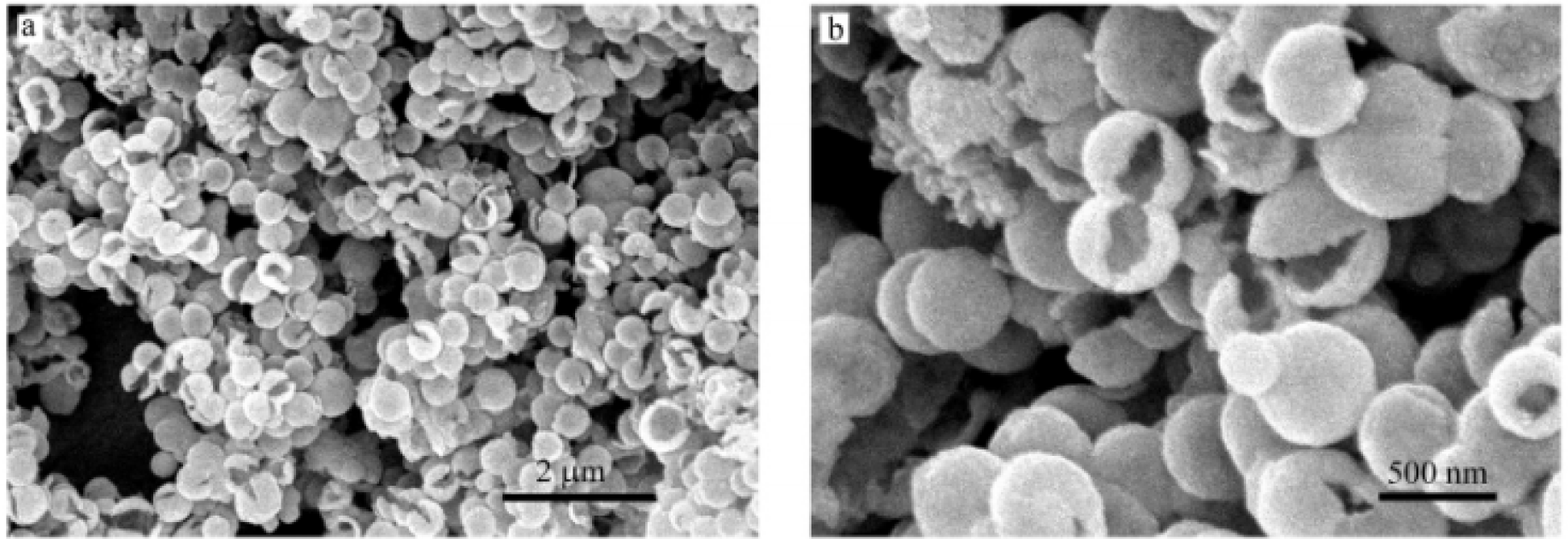
SEM analysis of carbon nanotubes.

### Electron Circulation

CPE, MWNT /Ag-TiO2 /CPE, MWNT /CPE, Ag-TiO2 /CPE electrodes were used as working electrodes at 1. 0 mmol / L 302 Analytical Chemistry Volume 38 [Fe(CN) 6]3 - /4 - + 0. 1 mol / L KCl mixed solution was characterized by cyclic voltammetry, the results are shown in Figure 2. Curve a is the cyclic voltammetry curve of [Fe(CN)6]3 - /4- on the bare CPE electrode. There is a relatively small number of redox peaks in the range of 1.2 to −0.8 V. At the MWNT /Ag-TiO2 /CPE electrode, a good peak of oxygen peak (curve b) with a significant increase in the peak height and a significant decrease in the redox peak potential difference was obtained, indicating that the MWNT /Ag-TiO2 modified electrode It has a larger surface area and better electrical conductivity. A pair of redox peaks were obtained on the MWNT /CPE electrode (curve c) and the Ag-TiO2 /CPE electrode (curve d), but their peak heights were significantly smaller than the curve b, and the peak potential difference was significantly larger than the curve b. MWNT and nano-Ag-TiO2 have strong synergistic effect on improving the surface electron conduction performance of the electrode. The reason may be: MWNT and nano-Ag-TiO2 can increase the active surface area; MWNT can be used as the super between Ag-TiO2 and the electrode surface. Micro-connector [20], which significantly improves the interface electron transfer capability.

**Figure 2.**
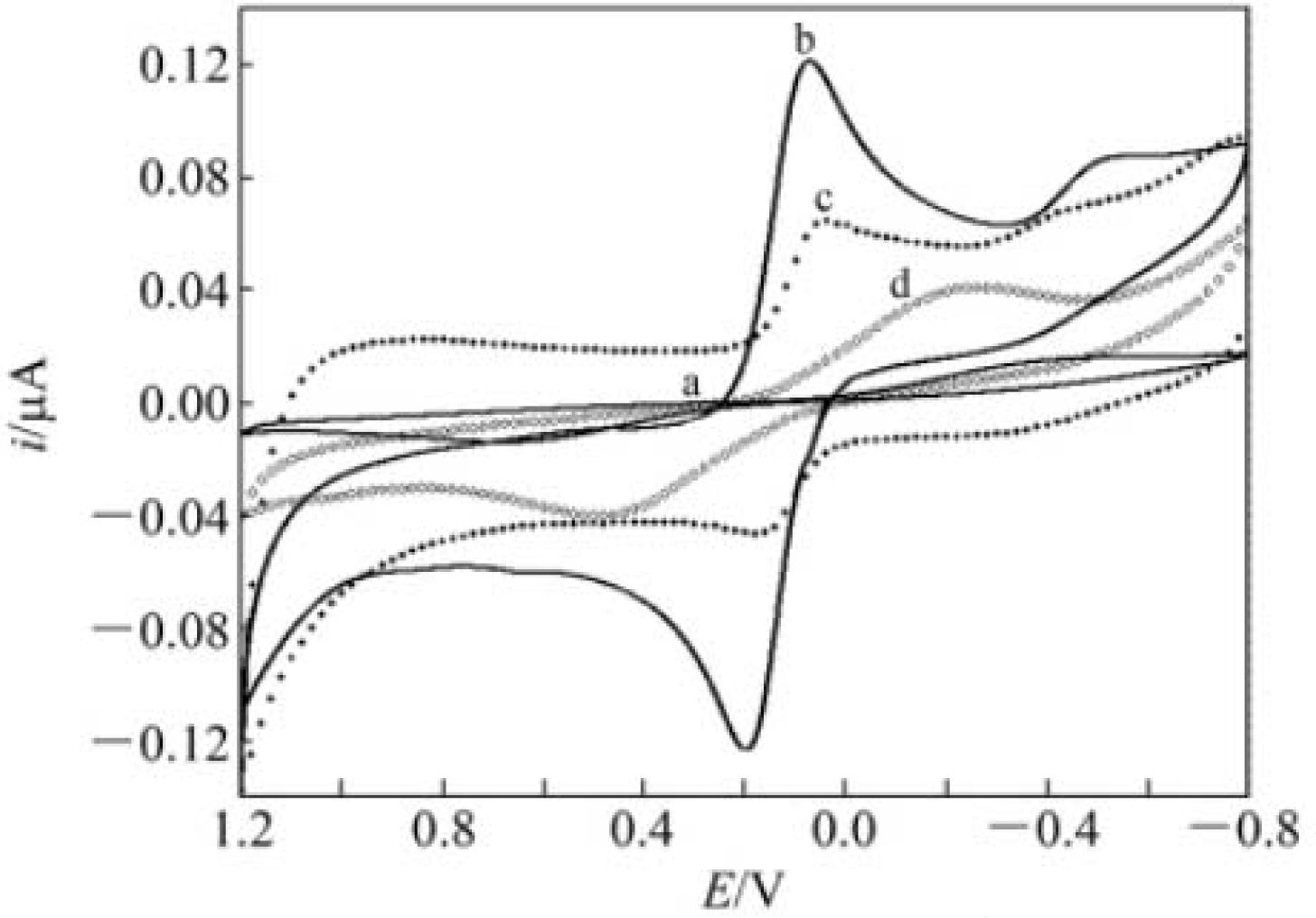
Electron migration chart.

### DNA immobilization

Figure 3 shows the electrochemical AC impedance Nyquist spectra of different working electrodes. On the electrode with MWNT /Ag-TiO2 modified film layer, the electron transfer resistance (Ret) of the interface is 520 Ω (curve b). Compared with curve a, the Ret of b is significantly reduced. This is consistent with the results of cyclic voltammetry. The Ag-TiO2 composite combines the advantages of both nano-TiO2 and Ag nanoparticles, exhibiting excellent biocompatibility and easy adsorption of biomolecules. There are a large number of defects and dangling bonds on the surface of MWNT, which are easy to be chemically modified. The modified MWNT can form a strong interaction with the nanocomposites. The ssDNA probe can be immobilized on the surface of the MWNT /Ag-TiO2 /CPE electrode due to the good electrical conductivity of the MWNT and the large specific surface area of the Ag-TiO2 nanocomposite, good electron transport properties and specific adsorption capacity for ssDNA.

**Figure 3.**
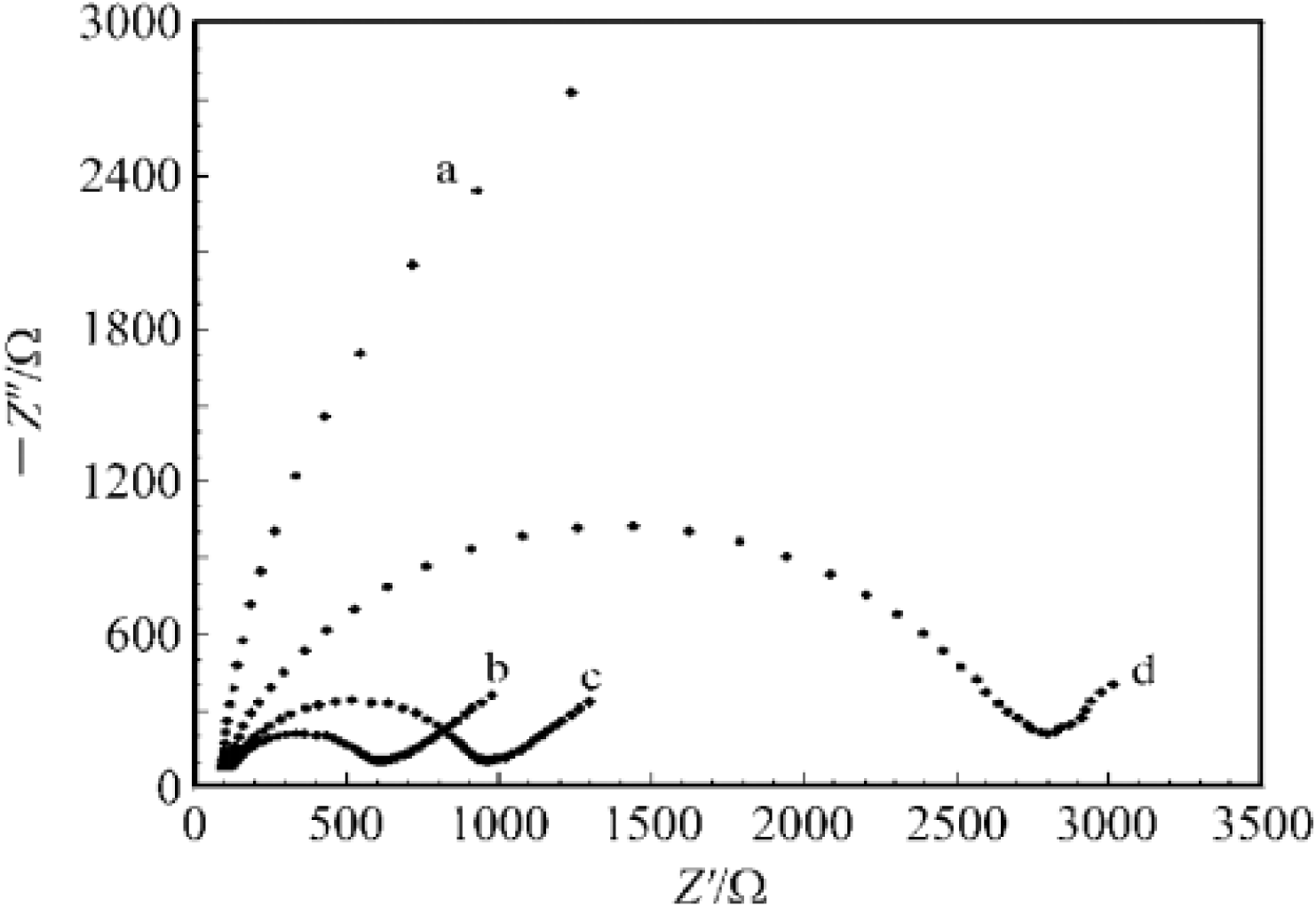
Electronic resistance analysis.

### Condition optimization

The fixed time directly affects the fixation of the DNA probe. Select 1. 0, 1. 5, 2. 0, 2. 5 and 3. 0 h as fixed time to detect the immobilization effect of the DNA probe. With the increase of the fixed time, the change of Ret on the surface of the probe DNA modification electrode and the change of Ret before fixation (ΔRet = RdsDNA - RssDNA) gradually increased. After 2.0 h, Ret and ΔRet were almost No longer change. In the experiment, 2 h was used as the probe DNA fixation time. Hybridization temperature has a large effect on hybridization efficiency and speed. Hybridization at 35 °C, 40, 45, 50 and 55 °C, recording AC impedance values, No. 303 3 Zhou Na et al: Multi-walled carbon nanotubes/nano Ag-TiO2 membrane DNA electrochemical biosensor Calculated ΔRet before and after hybridization. The results show that ΔRet increases with the increase of hybridization temperature between 35 and 45 °C. When the hybridization temperature continues to increase, the ΔRet value decreases slightly. The probe DNA was hybridized with the cDNA for 10 to 100 min at 45 °C, and the ΔRet value at each hybridization time was calculated. ΔRet increases with time between 10 and 60 min; as the hybridization time continues to increase, the ΔRet value no longer increases. Therefore, the optimal hybridization conditions were hybridization at 45 °C for 60 min.

### PAT gene sequencing

The dsDNA/MWNT /Ag-TiO2 /CPE formed by hybridization of the probe DNA with different DNA sequences at 0. 0 mmol / L K4Fe(CN) 6 /K3Fe(CN) 6 (1:1, V /V) The difference ΔRet of the electron transfer resistance values in the impedance spectrum of 1 mol / L KCl solution is used as a measurement signal to detect a 20-base PAT gene fragment. Figure 4 shows the histogram of ΔRet for different DNA sequences. The ΔRet measured by hybridization of probe DNA with non-complementary DNA (ncDNA) is very small. The ΔRe of the probe DNA hybridized with the 2 and 1 base mismatch DNA sequences increased to different extents. The ΔRet is maximized by hybridization of the probe DNA to the cDNA. Therefore, this sensor not only recognizes the target cDNA and ncDNA well, but also recognizes a DNA sequence with 2 base mismatches or even 1 base mismatch. The 20-base PAT gene fragment was detected by using ΔRet measured after hybridization with the target DNA, and the result is shown in Fig. 5. The logarithm of the concentration of the PAT gene fragment was plotted as the average of three parallel measurements. The results showed that the average value of ΔRet and the concentration of the PAT gene fragment was 1.0 × 10 - 11 ∼1. 0 × 10 - 6 mol / L The logarithmic value has a good linear relationship. The linear regression equation is ΔRet (Ω) = 342. 4lgC + 3969, γ = 0. 9970. The standard deviation of the blank solution for 11 times was σ. The detection limit of the PAT gene fragment was 3.12 × 10 - 12 mol / L by electrochemical impedance spectroscopy according to the 3σ method. It can be seen that the DNA electrochemical biosensor constructed under the optimized conditions of immobilization and hybridization has strong recognition ability for the sequence of target DNA.

**Figure 4.**
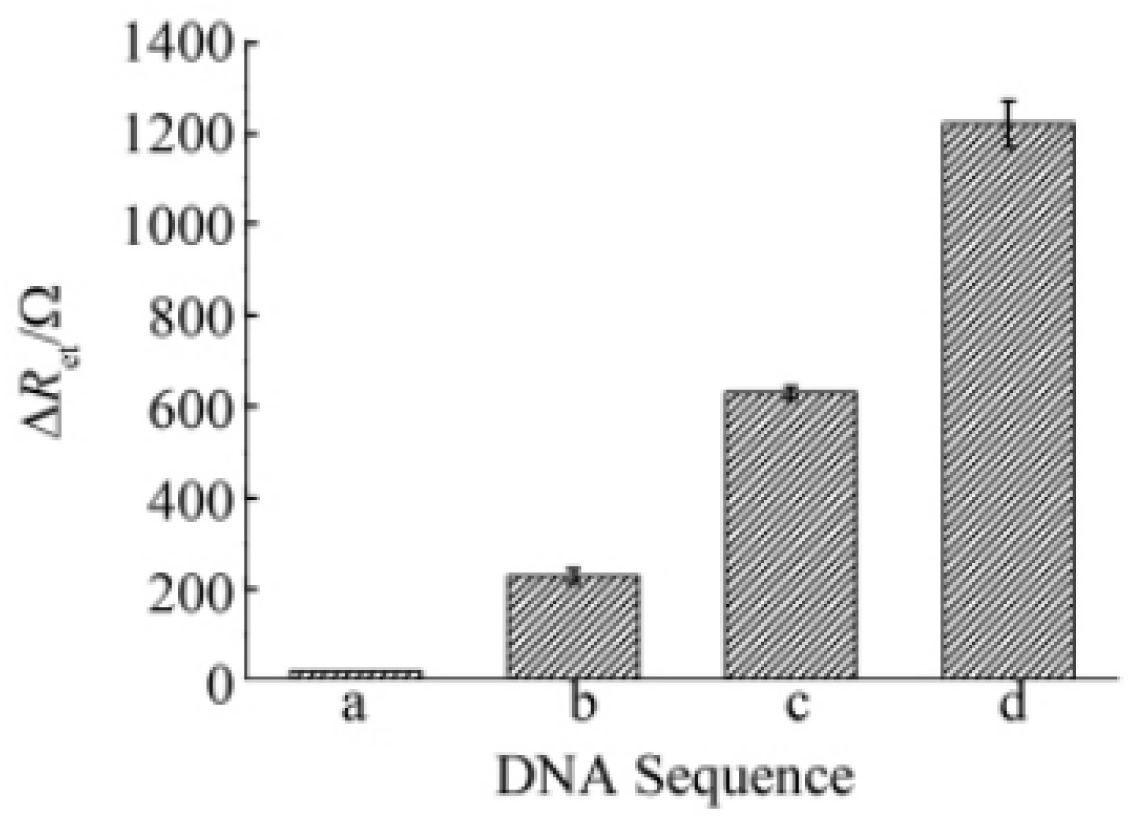
Detection method of ssDNA.

**Figure 5.**
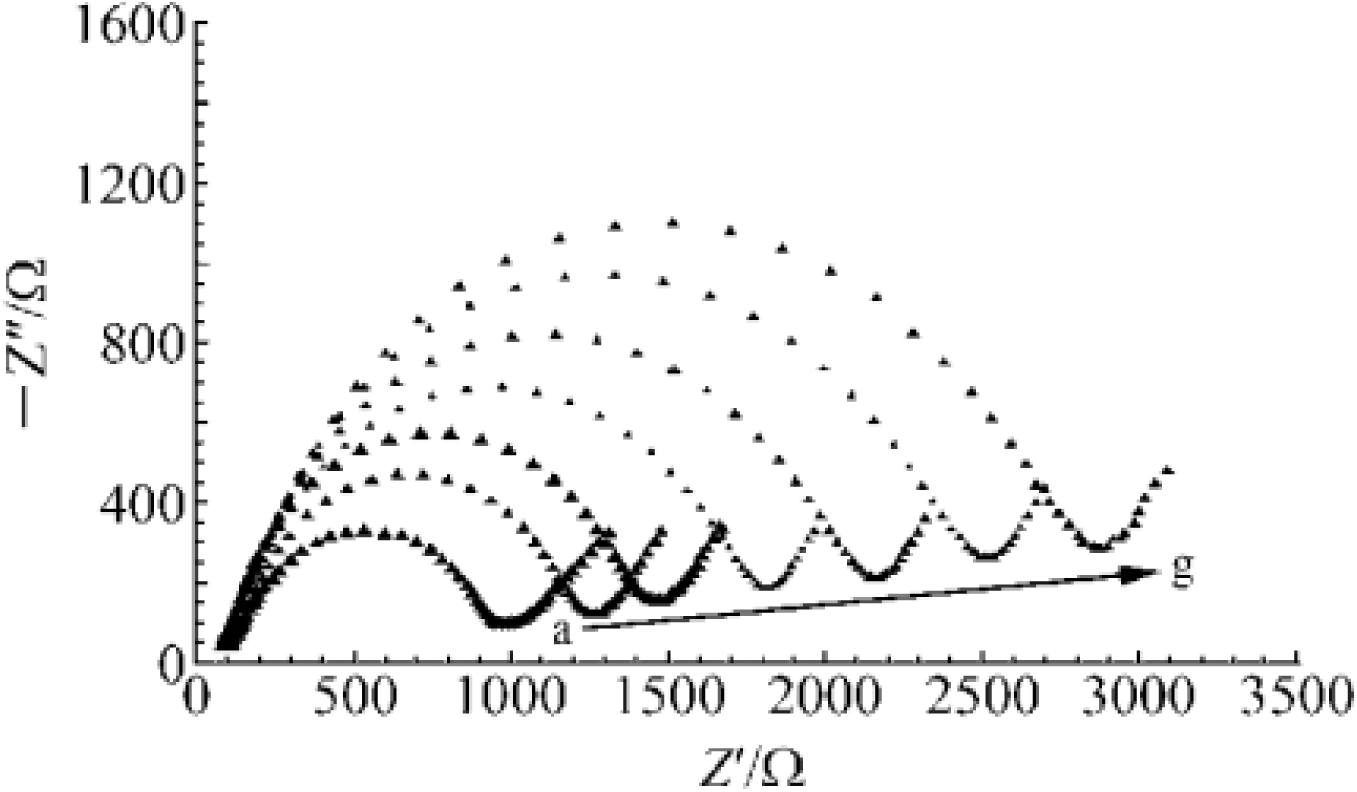
PAT analysis of gene expression correlates to resistance.

### DNA sensor repeatability assessment

The pole was placed in boiling water for 8 min, then quickly placed in an ice salt bath for 5 min to cool it. After washing, the electrochemical impedance spectroscopy spectra were recorded. The curve obtained was close to the probe electrode curve because of hybridization. The formed dsDNA is denaturing and depolarized at high temperature, and is converted into an ssDNA/MWNT /Ag-TiO2 /CPE electrode. After the heat-denatured ssDNA-modified electrode was hybridized again with the complementary DNA and subjected to the AC impedance test, the AC impedance value was again significantly increased, which was close to the impedance value of the dsDNA/MWNT/Ag-TiO2/CPE electrode obtained by the first hybridization. This sensor has better regenerative capacity. However, after 4 regenerations, the AC impedance value drops rapidly, possibly due to the ssDNA immobilized on the electrode. The reproducibility of this DNA electrochemical biosensor was investigated. Five electrodes were separately prepared under the same conditions, and the target DNA of 10. 0 × 304 analytical analysis No. 38 10 - 10 mol / L was determined, and the relative standard deviation of the five determinations was 4.32%. This sensor has good reproducibility. The sensor was stored at 4 °C for 20 d with no significant change in performance. Compared with the preparation of DNA electrochemical biosensors based on metal oxide composite nanomaterials reported in [21,22] (Table 1), the biosensor prepared by this method has lower detection limit, better reproducibility and reproducibility.

## Conclusion

A DNA electrochemical biosensor was prepared by using MWNT / Ag-TiO2 nanocomposite film. The composite membrane combines the advantages of MWNT, nano-TiO2 and Ag nanoparticles, and the synergistic effect between the nanoparticles greatly increases the immobilization of the DNA probe, thereby enabling highly sensitive detection of DNA hybridization. The preparation of MWNT/Ag-TiO2 nanocomposite film was followed by cyclic voltammetry. The immobilization and hybridization of DNA on the modified electrode were characterized by electrochemical impedance spectroscopy. The exogenous PAT gene fragment of transgenic maize was detected by electrochemical impedance spectroscopy. The sensor has good stability, reproducibility and regenerability.

